# Improved differential expression analysis of miRNA-seq data by modeling competition to be counted

**DOI:** 10.1101/2024.05.07.592964

**Authors:** Seong-Hwan Jun, Marc K. Halushka, Matthew N. McCall

## Abstract

MicroRNAs play a central role in regulating gene expression and modulating diseases. Despite the importance of microRNAs, statistical methods for analyzing them have received far less attention compared to messenger RNAs. Commonly, messenger RNA-seq methods are applied to microRNA-seq data, which may produce erroneous results due to the highly competitive nature of microRNA sequencing. This study critically examines and challenges the assumptions of messenger RNA-seq methods when applied to microRNA-seq data. We propose a Negative Binomial Softmax Regression (NBSR) method to model the unique characteristics of microRNA-seq data. On both simulated and experimental datasets, NBSR outperforms existing methods and offers a new perspective for analyzing microRNA-seq data. NBSR is implemented in Python and freely available as open-source software.

---

MicroRNA-seq (miRNA-seq) data analysis often utilizes methods developed for bulk messenger RNA-seq (mRNA-seq), such as DESeq2 (Love et al., 2014; Zhu et al., 2019) and edgeR (Robinson et al., 2010; Chen et al., 2024). A fundamental aspect of these methods is the approximate independence assumption, allowing each feature (e.g. mRNA or miRNA) to be analyzed independently. This assumption holds particularly well when there is a large number of unique RNA molecules, and the sequencing reads are fairly evenly distributed across these molecules. Under these conditions, the complex multivariate distribution of RNA molecules can be effectively approximated by a collection of univariate distributions (Townes et al., 2019).

Pertaining to miRNA-seq data, we postulate that there is non-negligible competition to be counted among microRNA (miRNA) species, where the increased sequencing reads from one miRNA necessarily decreased the observed sequencing read from another miRNA. In particular, the competition for sequencing significantly impacts lowly expressed miRNAs, where fold changes often appear large due to a small and noisy baseline. Consequently, the common practice of selecting significantly differentially-expressed miRNAs based on the absolute magnitude of the fold change exceeding a threshold may warrant scrutiny. For instance, Santangelo et al. (2016) filters miRNA by absolute FC > 1.5 and FDR < 0.1. Similarly, examining Table 1 of Satoh et al. (2015), the six differentially-expressed miRNAs with the largest positive fold changes have low baseline expression. This suggests that reliance on fold change thresholds without considering baseline abundance may lead to misleading interpretations.

To contextualize the compositional nature of miRNA-seq relative to mRNA-seq, first note the total number of miRNA species discovered in humans is approximately 500 to 2,000 (Griffiths-Jones et al., 2006; Fromm et al., 2022), which is 10-40 times less than the over 20,000 genes found in humans (Collado-Torres et al., 2017). Furthermore, the distribution of reads is often highly skewed towards a small number of highly expressed miRNAs. In examining the microRNAome dataset (McCall et al., 2017), we observed that fewer than 50 miRNAs often take up 90% of the reads (Figure 1 **A**). Examining bulk mRNA-seq samples in the recount2 dataset (Collado-Torres et al., 2017), we found that more than 5,000 mRNAs are needed to account for 90% of the total reads (Figure 1 **B**). The coverage for expressed miRNAs further deviates from uniformity and is skewed towards an even smaller number of miRNAs as measured by the Shannon diversity index (Figure 1 **C**), while coverage is fairly evenly spread across mRNAs (Figure 1 **D**). We also noted a disproportionate number of reads are attributed to the most highly expressed miRNAs, accounting for approximately 20% of the total reads (Figure 1 **E**). In contrast, the most highly expressed mRNAs typically only account for 1-2% of the total reads in bulk mRNA-seq data (Figure 1 **F**). Finally, the Spearman’s correlation between the two most highly expressed miRNAs was negative across a range of datasets (Figure 1 **G**), while no such pattern was observed for mRNA (Figure 1 **H**).

**Figure 1:**
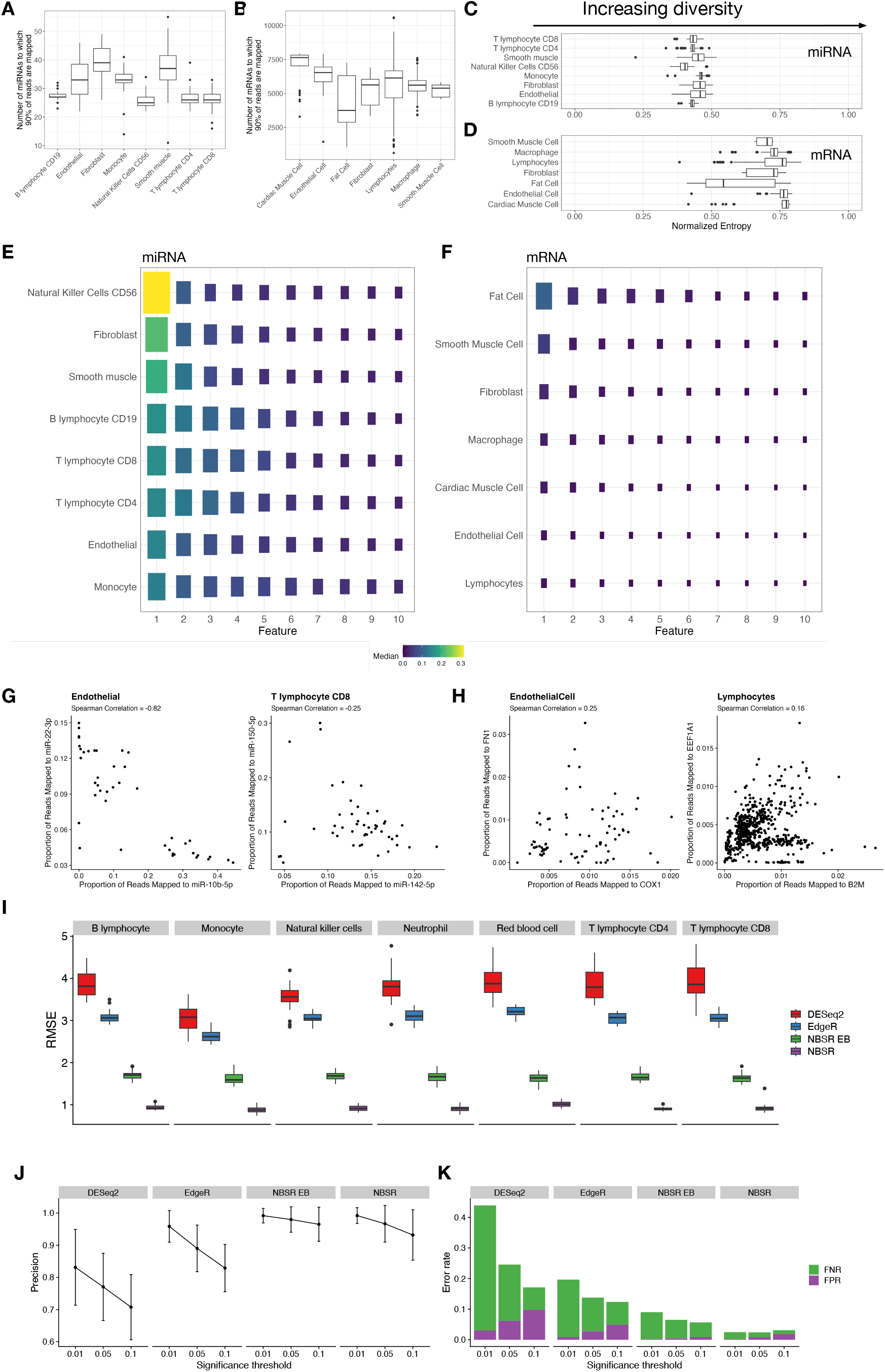
**A-B**. The number of features needed to account for 90% of the reads for miRNA and mRNA. **C-D**. The Shannon diversity measure for the miRNAs and mRNAs. **E-F**. Median of the proportion of reads allocated to the top 10 most highly expressed miRNAs and mRNAs. **G-H**. Spearman’s correlation for miRNA samples from Endothelial and CD8 T lymphocyte, and corresponding mRNA samples from Endothelial and Lymphocyte. **I-K**. Root mean squared error, precision, and error rates for DESeq2, EdgeR, NBSR EB, and NBSR for simulated data. The boxplots depict the interquartile range 1-3. The simulated data analysis was performed over 20 replicates. The confidence interval range shows ±2 standard errors from 20 replicates.

To address competition for expression, we propose a Negative Binomial Softmax Regression (NBSR) model, extending independent Negative Binomial regression by relating the underlying parameters using the softmax function. Specifically, the softmax regression models the fraction of all miRNA sequences in sample *i* originating from feature *j*, denoted *π*_*ij*_; this quantity represents the relative abundance level of *j* in sample *i* (Methods). The observed counts *Y*_*ij*_ are modeled using a Negative Binomial distribution, with the mean given by the product of its scale factor *s*_*i*_ and *π*_*ij*_. NBSR facilitates the modeling of dispersion parameters in relation to the relative abundance levels of miRNAs. The interpretation for the dispersion parameter is provided in terms of the biological coefficient of variation (CV), which measures the variability in *π*_*ij*_ in relation to its expectation between biological replicates (McCarthy et al., 2012; Chen et al., 2014). In contrast, mRNA-seq methods typically assume a shared dispersion parameter for each feature across all experimental conditions; this assumption may be violated in situations where there is a significant change in the miRNA expression across experimental conditions, altering the underlying distribution of the miRNA composition. Our approach allows for a more accurate depiction of biological variability, especially for miRNAs expressed at lower levels, which might be obscured by methods that model dispersion as solely depending on mean expression levels. By distinguishing the relative abundance represented by *π*_*ij*_ from the scale factor, our model aims to clarify the influence of each on observed expression variability.

We evaluated the NBSR on a simulated dataset where the approximate independence assumption is violated, comparing it to two widely-used mRNA-seq methods, DESeq2 and EdgeR, across two conditions (control vs. treatment) with *n* = 10 samples for each condition (Methods). In addition, we considered NBSR with Empirical Bayes estimation (NBSR EB), where we use NBSR to model competition for expression but assume a shared dispersion parameter for each miRNA (Methods). We computed the root mean squared error, confidence interval (CI) coverage, and precision for each of the methods. NBSR exhibited RMSE < 1 across different simulation settings (Figure 1 **I**). The desired nominal CI of 95% was attained (Supplemental Figure S2 **A**). We also compared the statistical power of the methods using false positive and negative rates; the Benjamini-Hochberg procedure was applied to the p-values obtained from each method at FDR thresholds of *α* ∈ {0.01, 0.05, 0.1}. NBSR achieved average precision > 0.9 across all FDR thresholds considered (Figure 1 **J**). NBSR is both stringent and sensitive as it achieved false positive rates of 0.201%, 0.879%, 1.87% and false negative rates of 2.29%, 1.54%, 1.25% for each of the thresholds considered, demonstrating the statistical power gained by modeling the relative abundances (Figure 1 **K**). We found that modeling the dispersion parameters in relation to the relative abundances further improved performance as NBSR outperformed NBSR EB in RMSE, coverage, and overall error rates. NBSR EB attained competitive precision and false positive rates to that of NBSR, demonstrating the performance gains from modeling competition for expression. Next, we derived the exact biological CV for the simulated data and compared it to the dispersion estimates obtained from the competing methods (Methods). As DESeq2, EdgeR, and NBSR EB assume one dispersion parameter per feature, these methods produced estimates that tend to fall between the true biological CVs for conditions A and B; NBSR estimates a range of dispersion values that align closely with the true biological CV (Figure 2 **A**). We also evaluated NBSR for different values of *n* and found that NBSR outperformed mRNA-seq methods in all measures considered (Supplementary Figures S2 **A-F**).

**Figure 2:**
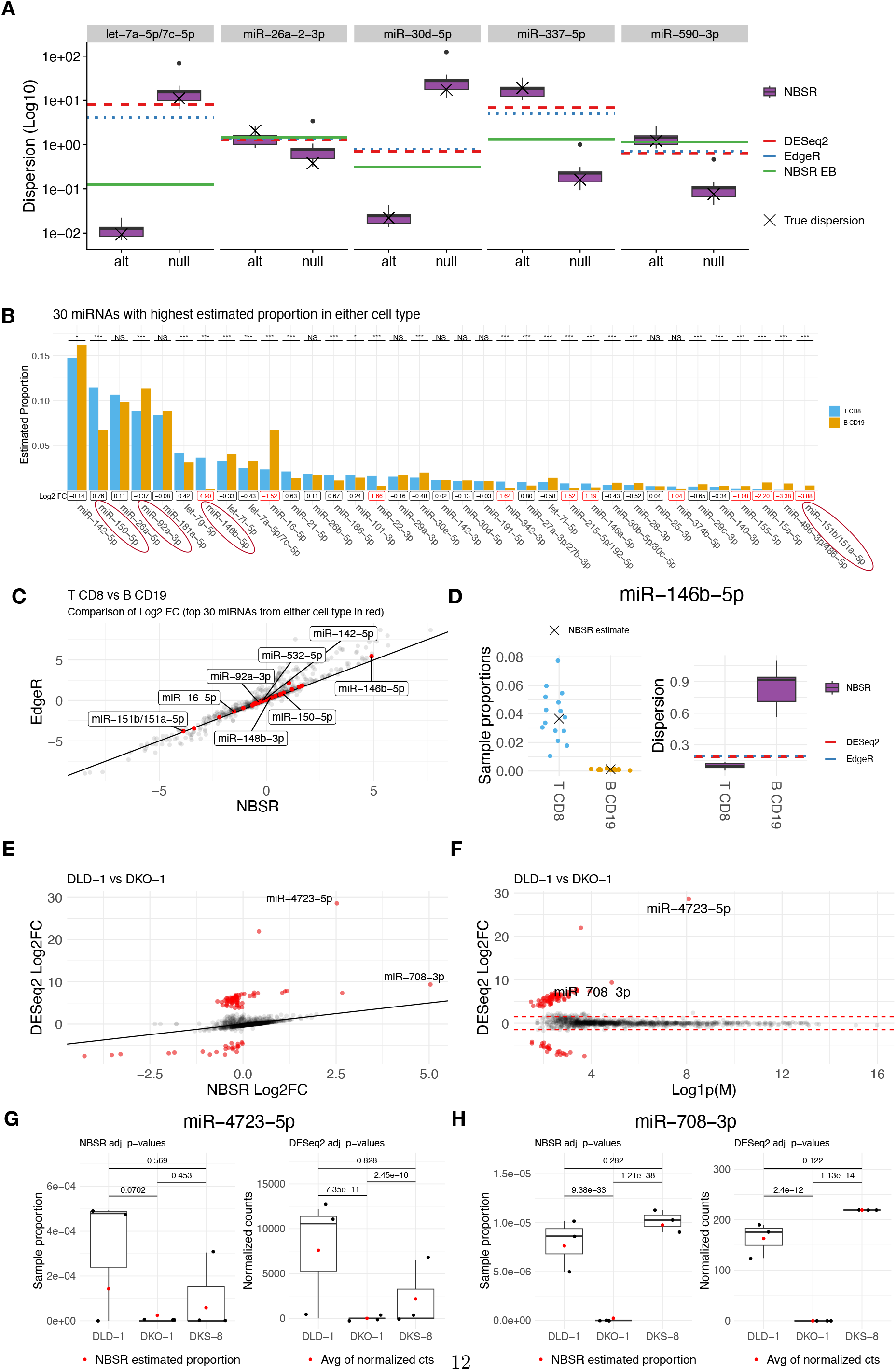
**A**. Dispersion estimates for a subset of the perturbed miRNAs from the simulated data analysis. The true biological CV (and dispersion) is marked by a cross for each miRNA. **B**. The 30 miRNAs with highest estimated relative abundance in either T (CD8+) or B (CD19+) cell types, resulting in 35 miRNAs in total across two cell types. Significance levels are denoted as *** at < 0.001, ** at < 0.01, and * at < 0.05. The fold change with T CD8 as the baseline is provided in log_2_ scale; miRNAs with at least a doubling in fold change are shown in red. **C**. Comparison of fold change estimates from EdgeR to NBSR for 35 miRNAs in **B. D**. In-depth examination of miR-146b-5p by visualizing the normalized counts from DESeq2 along with the dispersion estimates from NBSR. **E-F**. The fold change estimates in log_2_ scale for DESeq2 and NBSR and MA-plot for DESeq2; highlighted in red are miRNAs with |DESeq2 log_2_ FC− NBSR log_2_ FC| > 3. **G-H**. Plots of sample proportion and DESeq2’s normalized counts for miR-4723-5p and miR-708-3p.

We demonstrate the pitfalls of using thresholding to filter for significant differential expression of miRNAs. We analyzed T (CD8+) and B (CD19+) cells from the microRNAome dataset, which includes 15 samples for each cell type post-filtering (Methods). We identified the 30 miRNAs with the highest estimated proportions from each cell type, totaling 35 unique miRNAs. These accounted for 93.06% and 92.67% of the total miRNA proportions in T CD8 and B CD19 cells, respectively (Figure 2 **B**). The log_2_ fold change estimates from the NBSR model generally align with those from EdgeR and DESeq2 except for a handful of lowly expressed features (Figure 2 **C**, Supplementary Figure S3 **A-B**). Employing a threshold of FDR < 0.05 and absolute log_2_ fold change > 1 would lead us to overlook 16 miRNAs (Figure 2 **B**). Notably, miRNAs with lower estimated proportions in one of the two cell types, such as miR-146b-5p and miR-151b/151a-5p, displayed large absolute fold changes of 4.90 and −3.88, respectively. Furthermore, NBSR dispersion estimates for miR-146b-5p and miR-151b/151a-5p across the two cell types differed, indicating that there may be a change in biological CV (Figure 2 **D**, Supplementary Figure S3 **F**). Conversely, miRNAs like miR-150-5p and miR-92a-3p, which are highly expressed in both samples, showed smaller absolute log_2_ fold changes of 0.76 and 0.37, despite having considerable differences in estimated proportions of 4.70% and 2.55%, respectively (Supplementary Figure S3 **G-H**). The mRNA methods and NBSR disagreed on significance of a handful of miRNAs at FDR < 0.05 (Supplementary Figure S3 **B**). We further examined miR-148b-3p (NBSR-/mRNA+) and miR-532-5p/miR-142-5p (NBSR+/mRNA-) as most of the disagreements involved lowly express miRNAs (log(*M* + 1) < 5). The sample proportions and the NBSR estimated proportions did not show discernible differences for miR-148b-3p between T CD8 and B CD19 samples, with absolute difference of 0.0000651 in the NBSR estimated proportions (Supplementary Figure S3 **C**). For miR-532-5p and miR-142-5p, we noticed more visible differences in the sample proportions and the NBSR estimated proportions, with the absolute differences in the NBSR estimated proportions given by 0.0000393 and 0.0145 respectively (Supplementary Figure S3 **D-E**).

To assess the performance of NBSR in studies with small sample sizes, we analyzed colon adenocarcinoma cell lines DLD-1, DKO-1, and DKS-8, each with *n* = 3 from Shirasawa et al. (1993). These data exhibit competition for expression with top two miRNAs, miR-10a-5p and miR-21-5p, taking up nearly 60% of the reads (Supplementary Figure S4 **A**). Notably, the mRNA-seq methods estimate larger fold changes than NBSR (Figure 2 **E** and Supplementary Figures S4 **B-C**); analysis with an MA-plot shows that these miRNA with large fold changes are lowly expressed (Figure 2 **F**). In particular, miR-4723-5p exhibited wide range in DLD-1 samples, with two samples having counts over 10,000, while another sample did not express it. As DKO-1 samples did not express miR-4723-5p (Figure 2 **G**), it led DESeq2 to estimate a log_2_ FC close to 30 for this miRNA; NBSR produced a conservative log_2_ FC of approximately 2.5, finding no significant difference. This underscores NBSR’s conservativeness when data exhibits high variability and sparsity. In the absence of sparsity in DLD-1 samples, both DESeq2 and NBSR concurred on statistical significance of miR-708-3p (Figure 2 **H**). Examining the 10 miRNAs with highest proportion per cell line yielded 11 unique miRNAs (Supplementary Figure S4 **D**). Notably, miR-10a-5p showed significant increases in DKO-1 cell line, with an increase of 5.08% over DLD-1 and 3.92% over DKS-8 (Supplementary Figure S4 **C**). The reported absolute log_2_ FC (and FDR) values are 0.463 (0.555) for DESeq2, 0.484 (0.571) for EdgeR, and 0.192 (0.000448) for NBSR. While the statistical significances should be taken with caution in small-sample studies, NBSR performs multi-way comparisons and detects significant changes in expression of miR-10a-5p, let-7i-5p, and miR-200a/b-3p even when DESeq2 is unable to (Supplementary Figures S4 **E-H**). These miRNAs have been previously reported as having potential roles as biomarkers in colorectal cancer (Liu et al., 2017; Zheng et al., 2021; Ma et al., 2021; Klicka et al., 2022). NBSR estimated proportions generally concur with those from mRNA-seq methods for moderately expressed miRNAs (Supplementary Figures S3 **A** and S4 **I**).

Overall, NBSR offers several desirable properties: (1) differential expression across experimental conditions has an intuitive interpretation as the change in the probability of expression using either the ratio (log relative risk) or the difference in relative abundances; (2) it does not require filtering for miRNAs, as sparsely expressed miRNAs are properly accounted for in relation to other miRNAs; (3) the relationship between the biological coefficient of variation and the relative abundance level of the feature can be directly modeled, increasing statistical power while achieving a tighter confidence interval compared to mRNA-seq methods; (4) enhanced sensitivity in detecting differential expression in both sparsely and highly abundant miRNAs. We anticipate that as single-cell miRNA-seq becomes increasingly feasible and widely-used, these benefits will be even more pronounced.

NBSR and its associated methodologies introduced in this paper provide a new perspective for viewing and analyzing miRNAs. There are two extensions to this model that would likely further improve performance: (i) considering positive correlation between miRNAs sharing the same precursor miRNA transcript, and (ii) considering different isoforms of miRNAs, called isomiRs.

## Methods

### microRNAome and recount2

For the dataset used in creating miRNA plots in Figure 1, we took cell types from the microRNAome dataset with ≥40 samples after eliminating samples with less than 100,000 reads. The table of cell types and the number of samples are shown in Supplementary Table 1. For the corresponding mRNA plots in Figure 1, we first removed any duplicated samples. We then removed samples whose average read length exceeded 200. Cell types with at least 10 samples were selected for analysis. The table of cell types and the number of samples are shown in Supplementary Table 2. To simulate data under different parameter settings, we estimated the parameters using cell type data originating from Juzenas et al. (2017) (study id 89 in microRNAome dataset) with at least 20 samples in the microRNAome datasets (Supplementary Table 3). For each cell type dataset, we selected human miRNAs in the MirGeneDB database (Fromm et al., 2022), which contains 596 definitive miRNAs, then filtered out samples with less than 100, 000 miRNA reads. The colon adenocarcinoma cell lines originate from Cha et al. (2015) (study id 17 in microRNAome dataset).

### Negative Binomial Softmax Regression

For each sample *i* = 1, *…, N*, we have a *P* -dimensional covariate vector *x*_*i*_ and observe counts *Y*_*ij*_ for miRNA *j* = 1, *…, K*. The observed counts are modeled using the Negative Binomial distribution,

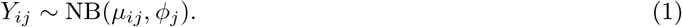

In mRNA-seq methods, the mean is modeled as *μ*_*ij*_ = *s*_*i*_*q*_*ij*_ with *s*_*i*_ denoting the scale factor for sample *i* and 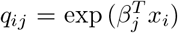. The effect of covariate *x*_*i*_ on miRNA *j* is determined by coefficients *β*_*j*_ ∈ ℝ^*P*^. The dispersion parameters *ϕ*_*j*_ accounts for overdispersion of the observed counts data.

To account for competition for expression, we propose to model the mean by *μ*_*ij*_ = *s*_*i*_*π*_*ij*_ where *π*_*ij*_ ∈ [0, 1] such that Σ_*j*_ *π*_*ij*_ = 1. We model *π*_*ij*_ using softmax regression:

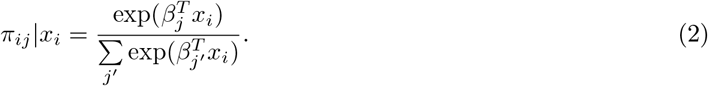

We denote *s*_*i*_ as the total number of counts: *s*_*i*_ = Σ_*j*_ *Y*_*ij*_. We place a Normal prior on coefficients *β*_*jp*_, centered around 0 with a standard error *σ*_*p*_ for each covariate *p* = 1, *…, P*. The Normal prior serves to regularize parameter estimation towards zero (Love et al., 2014). The complete model is specified as follows,

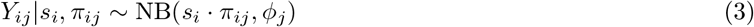

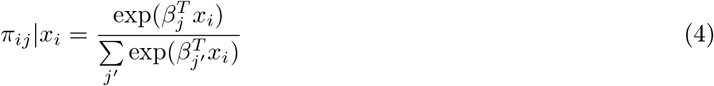

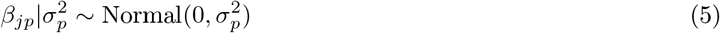

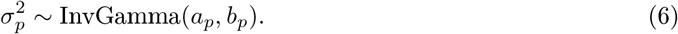

The parameters of the Inverse Gamma are chosen by the user; we found values of *a*_*p*_ = 3, *b*_*p*_ = 2 to work well in many situations and suggest these as reasonable defaults.

### Biological coefficient of variation

We examine the unconditional coefficient of variation (CV) over *Y*_*ij*_, as considered in McCarthy et al. (2012) and Chen et al. (2014), and show that it motivates the need for re-modeling dispersion. As above, let *π*_*ij*_ ≥ 0 be a random variable denoting the fraction of feature *j* in sample *i* such that Σ_*j*_ *π*_*ij*_ = 1. Let us denote the CV for *π*_*ij*_ by 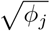. By denoting 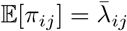 and assuming that the CV is fixed for *j* across biological samples *i*, we necessarily obtain that 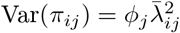 from the definition of CV. The unconditional CV of *Y*_*ij*_ can be decomposed into technical and biological CV as follows:

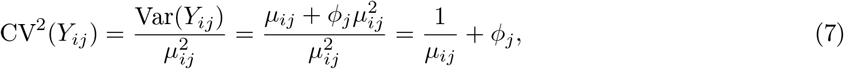

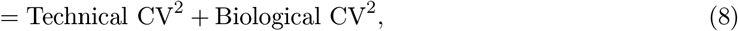

where Var(*Y*_*ij*_) can be obtained from law of total variance,

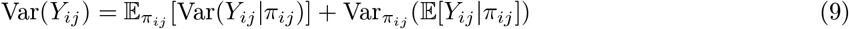

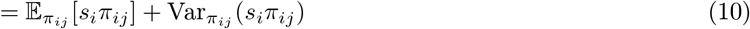

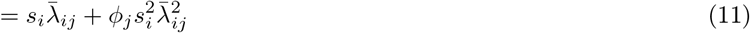

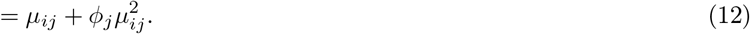

This decomposition states that the technical CV stems from variability in *Y*_*ij*_ due to experimental settings for a fixed sample *i* (e.g., variability from sequencing equipment, library preparation method, and other experimental factors). This technical variability is captured using a Poisson distribution and as such, 1*/μ*_*ij*_ is the square of the coefficient of variation of a Poisson distribution with a mean *μ*_*ij*_. On the other hand, *ϕ*_*j*_ captures the biological variability in *π*_*ij*_ across different biological replicates. This justifies the need for modeling dispersion in terms of the relative abundance *π*_*ij*_, which is only possible if we use a model that directly represents *π*_*ij*_. The second observation concerns the assumption of a fixed CV across biological samples for *j*, from which the common practice of assuming a shared dispersion parameter for each feature in mRNA-seq methods stems. However, as we typically consider two or more experimental conditions, the underlying distribution and moments of *π*_*ij*_ may significantly differ across experimental conditions, violating this assumption.

The above decomposition further suggests that when the mean expression is high, much of the variation is of a biological nature, while when the mean is low, the technical variability can dominate. In the case of miRNA, the technical variability can exacerbate the difficulties associated with dispersion estimation. Specifically, miRNA-seq data is usually obtained using small RNA sequencing (sRNA-seq) technology, where an additional source of technical variability stems from the total abundance of miRNAs in relation to other small RNA molecules targeted by sRNA-seq in a given sample and the miRNA capture rate of an experiment (Lu et al., 2018). Indeed, we noted that there was minimal correlation between the percentage of reads mapping to miRNAs (percent miRNA) and the total input reads (Supplementary Figure S1 **A**). While the total sum of miRNA reads is highly correlated with the percent miRNA (Pearson’s correlation of 0.567), there remains significant technical variability in the miRNA capture rate across samples leading to variability in total miRNA reads for the same percent miRNA (Supplementary Figure S1 **B**). These technical variabilities can inflate dispersion estimation as seen in Supplementary Figure S1 **C-D**, where large dispersion is estimated for lowly expressed and highly sparse miRNAs for the immune cell data analyzed using DESeq2. Supplementary Figure S1 **E** shows that mRNA-seq methods estimate large dispersion to account for zero expression (sparsity): the samples whose observed (normalized) counts are zero (x-axis) tend to have relatively large predicted counts are necessarily going to be over-dispersed. For these lowly expressed features, the normalized abundance *π*_*ij*_ may be more informative than the mean expression *μ*_*ij*_ because the true relationship between the dispersion and mean may be obscured by the size factor estimation.

### Modeling dispersion

The current practice employed in mRNA-seq methods is to model dispersion as a function of the mean expression level of the feature (Love et al., 2014; Chen et al., 2014; Eling et al., 2018; Risso et al., 2018). In the original DESeq2 paper, the relationship between the dispersion and mean expression is modeled using a log-normal distribution centered around a trend line specified as a function of the mean of normalized read counts. EdgeR deploys a similar strategy by decomposing dispersion into three components, global level dispersion, trended dispersion, and tagwise (feature-specific) dispersion. These strategies have been adopted in the scRNA-seq domain with modifications, for example, using a t-distribution and a semi-parametric model for robustness and flexibility (e.g., Eling et al. (2018)). In all of these approaches, the dispersion is modeled in relation to the feature-specific mean expression levels *μ*_*j*_ or the mean of suitably normalized counts 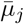, both quantities are affected by the scale factor of the samples. However, as we noted above, it may be beneficial to decouple dispersion from the scale factor as the dispersion is a quantity that pertains to *π*_*ij*_. We propose to model the dispersion in relation to *π*_*ij*_ and the total miRNA reads *R*_*i*_ =Σ _*j*_ *Y*_*ij*_:

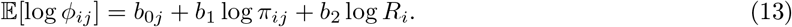

In essence, we include *R*_*i*_ to model technical variability such as the miRNA capture rate. By considering *π*_*ij*_ and *R*_*i*_ as separate input variables, we are able to decouple the effect of the scale factor from the relative abundance *π*_*ij*_. While a full Bayesian treatment would involve specifying a prior over log*ϕ*_*ij*_ with the expectation given in Equation 13, our implementation opts for a point estimation of *ϕ*_*ij*_ by optimizing parameters *b*_0*j*_, *b*_1_, *b*_2_.

In our simulated study, with *n* ∈ {10, 20}, the above dispersion model successfully captured the true biological CV (Figure 2 **A** and Supplementary Figure S2 **G**). However, for *n* ∈ {3, 5}, we observed convergence issues with the optimization, likely due to an insufficient number of data points leading to an unidentifiable posterior distribution. For studies with a small number of samples, we therefore recommend assuming a fixed biological CV across samples (i.e., feature-specific dispersion assumption) and estimating dispersion using NBSR EB, a multi-stage empirical Bayes (EB) procedure as adopted by DESeq2 and EdgeR. Namely, we first estimate and fix the mean expression 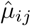 using DESeq2 (or EdgeR) and obtain the MLE 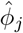. We then elicit a prior distribution over the dispersion for each feature by regressing 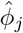 on 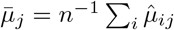 using local regression with a Gaussian Radial Basis Function (GRBF) to obtain the mean function (Eling et al., 2018).

The NBSR EB model is given with a prior on dispersion as follows:

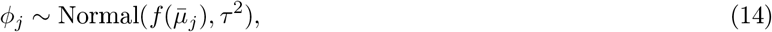

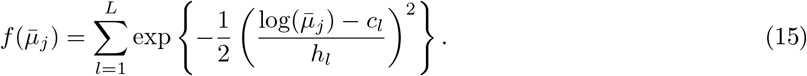

We used *τ* = 0.5 in the experiments. We set the default values for the GRBF following recommendations from Eling et al. (2018), with *L* = 10 and equi-spaced centers *c*_*l*_ from min_*j*_ 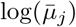 to max_*j*_ 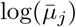, and *h*_*l*_ = 1.2 · (*c*_2_ − *c*_1_). In the colon adenocarcinoma experiments, we used NBSR EB with the above default values. For analysis of simulated data for *n* ∈ {3, 5}, we used NBSR EB but rather than using GRBF prior over *ϕ*_*j*_, we fixed the dispersion at the mean 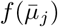 and focused on quantifying performance of NBSR (Supplementary Figures S2 **A-F**).

### Inference on fold change

We are typically interested in comparing two experimental conditions, A and B. Without loss of generality, we take A as the baseline level. We aim to test the null hypothesis that there is no difference in the expression levels across the two conditions against the two-sided hypothesis that the expression is different. The mRNA-seq methods carry out hypothesis testing directly on *β*_*j*_ to determine the significance of feature *j* on the outcome. NBSR aims at detecting direct changes to the expression level of feature *j* and indirect changes due to changes in other features *j*′ ≠ *j*. To measure this indirect effect, we compare the conditional probability for condition B in relation to condition A, denoted *π*_*j*|*B*_ and *π*_*j*|*A*_ respectively. One possibility is to consider the log ratio,

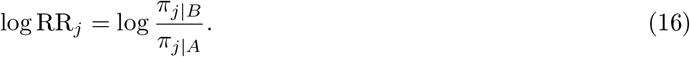

This quantity has a nice interpretation in statistics as the log of *relative risk* as *π*_*j*|*B*_ represents the probability of an outcome for group B and similarly for *π*_*j*|*A*_. We developed an inference methodology based on the log relative risk in relation to the parameters of the NBSR. We derive the closed form for the Hessian matrix of the coefficients *β* and use the pseudo-inverse to obtain the Fisher information matrix. We found the pseudo-inverse to be fast and accurate even when the exact inverse was not available. Statistical inference on the differential expression of each miRNA utilizes the log of the relative risk and a complete derivation of the standard error estimates of log_2_ fold change is provided in the Supplementary Information.

### Simulated data generation

We use the Dirichlet-Multinomial distribution or a generalized Polya’s urn experiment to simulate competition to be counted (see for e.g., Koptagel et al. (2024)). The Dirichlet model necessarily induces a negative correlation between the features, which makes it well-suited for simulating competition to be counted. We sample the probability vectors ***π***_*i*_ from a Dirichlet distribution with parameters ***λ*** = (*λ*_1_, *…, λ*_*J*_) ≥ **0**. We sample the sequencing depth *d*_*i*_∼ Uniform(10^6^, 10^8^), sample the miRNA capture rate *z*_*i*_∼ Uniform(0, 0.5) independently, and set the miRNA reads *s*_*i*_ = ⌈*z*_*i*_ · *d*_*i*_⌉. The read counts *Y*_*ij*_ are sampled from a Multinomial distribution with ***π***_*i*_ and *s*_*i*_ as parameters.

We obtained maximum likelihood estimates of ***λ*** for each cell type data (Supplementary Table 3) using the observed sample proportions *p*_*ij*_ = *Y*_*ij*_/ Σ _*j*_ *Y*_*ij*_. The mean of the Dirichlet distribution given by 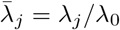 where *λ*_0_ = Σ_*j′*_ *λ*_*j′*_ is referred to as the precision parameter. We estimate *λ*_*j*_, *λ*_0_ using the fixed-point iteration described in Minka (2000). Additional details of the estimation procedure are presented in the Supplementary Information.

To simulate an experimental set up, we set the parameter values for the Dirichlet distribution representing the control group as 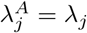. We then classify the miRNAs for each dataset into two classes: *𝒥*_*LOW*_ = {*j* : *λ*_*j*_ ≤ 1} and *𝒥*_*HIGH*_ = {*j* : *λ*_*j*_ > 1}. We sample *K* = 40 pairs of indices 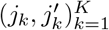 such that *j*_*k*_ ∈ 𝒥_*LOW*_ and 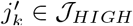 and swap their parameters to form the ground truth parameters for the treatment condition *B*, i.e., 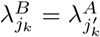 and 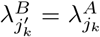. The ground truth log_2_ fold change for miRNA *j* is given by 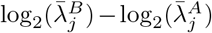.

The read counts are then simulated from a Multinomial distribution: 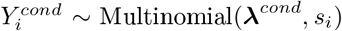 for *cond* ∈ {*A, B*}.

As we sample ***π***_*i*_ ∼ Dirichlet(*λ*_1_, …, *λ*_*J*_), we can derive the exact biological CV as follows:

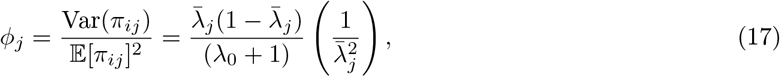

where *λ*_0_ = Σ _*j′*_ *λ*_*j′*_ and 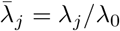. We can compute the ground truth dispersion for each condition A and B using Equation 17. In such a case, the identical distribution assumption is no longer true and *π*_*i*|*A*_, *π*_*i*|*B*_ arise from Dirichlet distributions.

## Data availability

The miRNA-seq data used in the analysis is available from `microRNAome` package through Bioconductor https://www.bioconductor.org/packages/3.16/data/experiment/html/microRNAome.html v1.20.0. The mRNA-seq data used in the analysis is available from recount package through Bioconductor https://bioconductor.org/packages/release/bioc/html/recount.html accessed on 01/06/2020. The cell types and the T CD8 vs B CD19 analysis data are available on sequencing read archive with accession SRP110505 under PRJNA391912. The colon adenocarcinoma cell lines are available on sequencing read archive with accession SRP056261 under PRJNA278673. The specific run accessions are SRR1917323, SRR1917324, SRR1917325 for DKO-1, SRR1917335, SRR1917336, (DLD-1), and SRR1917337, SRR1917329,SRR1917330, SRR1917331 (DKS-8).

## Code availability

We implemented NBSR model using PyTorch and utilize automatic gradient computation to optimize the model parameters (Paszke et al., 2019). The code is available on Github https://github.com/junseonghwan/nbsr/.

