## Supplementary Materials for "Improved differential expression analysis of miRNA-seq data by modeling competition to be counted"

Seong-Hwan Jun[1], Marc K. Halushka[2], and Matthew N. McCall[1]

[1]Department of Biostatistics and Computational Biology, University of Rochester Medical Center
[2]Pathology and Laboratory Medicine Institute, Cleveland Clinic

#### 1 Laplace's approximation

We derive standard error estimates via Laplace's approximation of the posterior distribution over the covariate coefficients $\beta = (\beta_{j,d})_{j,d}$ for $j = 1, ..., K$ and $d = 1, ..., P$, evaluated at the MAP estimate given by, Normal$(\hat{\beta}_{MAP}, \hat{\Sigma})$, see e.g., (Gelman et al., 2013, Section 13.3). Typically, the covariance matrix is estimated by the inverse of the observed information, which is given by the negative of the Hessian of the log posterior distribution evaluated at the MAP estimate.

#### 2 Gradients for NBSR parameters

We begin by deriving the gradient of the log posterior distribution, which is given by

$$\log p(\beta|Y) \propto \log p(Y|\beta) + \log p(\beta).$$

We derive the first order gradient of the log likelihood of the NBSR model with respect to $\beta_{k,d}$, which we denote $\nabla_{k,d} \log p(Y|\beta)$.

$$\nabla_{k,d} \log p(Y|\beta) = \sum_i \sum_j \nabla_{k,d} \log p(y_{i,j}|\beta). \tag{1}$$

Note that $p(y_{i,j}|\beta)$ is given by Negative Binomial distribution with parameters $p_{i,j} = \mu_{i,j}/\sigma^2_{i,j}$ and $r_j = 1/\phi_j$:

$$\log p(y_{i,j}|\beta) = \log C_{i,j} + r_j \log p_{i,j} + y_{i,j} \log(1 - p_{i,j}), \tag{2}$$

where $C_{i,j}$ is a constant with respect to $\beta$. We will further simplify the expression by plugging-in $p_{i,j} = \mu_{i,j}/\sigma^2_{i,j}$:

$$\log p(y_{i,j}|\beta) = r_j(\log \mu_{i,j} - \log \sigma^2_{i,j}) + y_{i,j}(\log(\mu^2_{i,j}) - \log \sigma^2_{i,j}). \tag{3}$$

Since $\sigma^2_{i,j} = \mu_{i,j} + \phi_j \mu^2_{i,j}$, computing the gradient amounts to computing the gradient of $\mu_{i,j}$ w.r.t $\beta_{k,p}$:

$$\begin{align} \nabla_{k,d}\mu_{i,j} &= s_i \nabla_{k,d}\pi_{i,j} \tag{4} \\ &= x_{i,d}\mu_{i,j}(1[j = k] - \pi_{i,k}), \tag{5} \end{align}$$

where the gradient for $\pi_{i,j}$ is given by,

$$\begin{align} \nabla_{k,d}\pi_{i,j} &= \nabla_{k,d} \frac{\exp(\beta_j^T x_i)}{\sum_{j'} \exp(\beta_{j'}^T x_i)} \tag{6} \\ &= \begin{cases} x_{i,d}\pi_{i,j}(1 - \pi_{i,k}) & \text{if } j = k \\ -x_{i,d}\pi_{i,j}\pi_{i,k} & \text{if } j \neq k \end{cases} \tag{7} \\ &= x_{i,d}\pi_{i,j}(1[j = k] - \pi_{i,k}). \tag{8} \end{align}$$

The gradient of $\mu_{i,j}^2$ can be derived by applying the chain rule:

$$\nabla_{k,d}\mu_{i,j}^2 = 2\mu_{i,j}\nabla_{k,d}\mu_{i,j} \tag{9}$$

$$= 2x_{i,d}\mu_{i,j}^2(1[j = k] - \pi_{i,k}). \tag{10}$$

Finally, the gradient of $\sigma_{i,j}^2$:

$$\nabla_{k,d}\sigma_{i,j}^2 = \nabla_{k,d}\mu_{i,j} + \phi_j\nabla_{k,d}\mu_{i,j}^2 \tag{11}$$

$$= x_{i,d}\mu_{i,j}(1[j = k] - \pi_{i,k})(1 + 2\phi_j\mu_{i,j}). \tag{12}$$

Putting all of this together, we have the following expression for the gradient of log likelihood:

$$\nabla_{k,d}\log p(y_{i,j}|\beta) = r_j x_{i,d}(1[j = k] - \pi_{i,k})\left(1 - \frac{\mu_{i,j}(1 + 2\phi_j\mu_{i,j})}{\sigma_{i,j}^2}\right) \tag{13}$$

$$+ y_{i,j}x_{i,d}(1[j = k] - \pi_{i,k})\left(2 - \frac{\mu_{i,j}(1 + 2\phi_j\mu_{i,j})}{\sigma_{i,j}^2}\right) \tag{14}$$

The gradient of $\log p(\beta)$ is,

$$\nabla_{k,d}\log p(\beta) = \nabla_{k,d}\log p(\beta_{k,d})$$

$$= -\frac{\beta_{k,d}}{\sigma_d^2}.$$

Note that

$$\mu_{i,j}(1 + 2\phi_j\mu_{i,j}) = \sigma_{i,j}^2 + \phi_j\mu_{i,j}^2,$$

allowing us to simplify Eq 13 further:

$$\nabla_{k,d}\log p(y_{i,j}|\beta) = r_j x_{i,d}(1[j = k] - \pi_{i,k})\left(-\frac{\phi_j\mu_{i,j}^2}{\sigma_{i,j}^2}\right) \tag{15}$$

$$+ y_{i,j}x_{i,d}(1[j = k] - \pi_{i,k})\left(1 - \frac{\phi_j\mu_{i,j}^2}{\sigma_{i,j}^2}\right) \tag{16}$$

Hence, the gradient of the log posterior distribution is given by,

$$\nabla_{k,d}\log p(\beta|Y) = \sum_i\sum_j -x_{i,d}r_j(1[j = k] - \pi_{i,k})D_{i,j} + y_{i,j}x_{i,d}(1[j = k] - \pi_{i,k})(1 - D_{i,j}) - \frac{\beta_{k,d}}{\sigma_d^2}, \tag{17}$$

14 where $D_{i,j} = \frac{\phi_j\mu_{i,j}^2}{\sigma_{i,j}^2}$.

We now derive the second derivative of the log posterior.

$$\nabla_{k',d'}\nabla_{k,d}\log p(y_{i,j}|\beta) = -x_{i,d}r_j\left[-\nabla_{k',d'}\pi_{i,k}D_{i,j} + (1[j = k] - \pi_{i,k})\nabla_{k',d'}D_{i,j}\right]$$
$$+ x_{i,d}y_{i,j}\left[-\nabla_{k',d'}\pi_{i,k}(1 - D_{i,j}) + (1[j = k] - \pi_{i,k})(-\nabla_{k',d'}D_{i,j})\right].$$

This amounts to computing the gradient of $D_{i,j}$ as we have derived the gradient for $\pi_{i,j}$ in Eq (6). The gradient of $D_{i,j}$ can be obtained by using a trick. For any function $f$

$$f(x)\nabla_x\log f(x) = \nabla_x f(x), \tag{18}$$

combined with the gradient of $\log\mu_{i,j}^2/\sigma_{i,j}^2$ we have in Eq (3), the gradient can be obtained as,

$$\nabla_{k,d}D_{i,j} = x_{i,d}(1[j = k] - \pi_{i,k})D_{i,j}(1 - D_{i,j}). \tag{19}$$

The Hessian for the log likelihood is given by,

$$
\begin{aligned}
H_{loglik} = \nabla_{k',d'} \nabla_{k,d} \log p(y_{i,j}|\beta) =& r_j D_{i,j} x_{i,d} \pi_{i,k} x_{i,d'} (1[k = k'] - \pi_{i,k'}) && (20)\\
&- y_{i,j}(1 - D_{i,j}) x_{i,d} \pi_{i,k} x_{i,d'} (1[k = k'] - \pi_{i,k'}) && (21)\\
&- r_j D_{i,j}(1 - D_{i,j}) x_{i,d}(1[j = k] - \pi_{i,k}) x_{i,d'} (1[j = k'] - \pi_{i,k'}) && (22)\\
&- y_{i,j} D_{i,j}(1 - D_{i,j}) x_{i,d}(1[j = k] - \pi_{i,k}) x_{i,d'} (1[j = k'] - \pi_{i,k'}). && (23)
\end{aligned}
$$

And the Hessian for the log posterior is given by,

$$H = H_{loglik} - \frac{1}{\sigma_d^2} I_{JP \times JP}. \tag{24}$$

Finally, we set the covariance matrix as,

$$\hat{\Sigma} = (-H)^{-1}. \tag{25}$$

### 3 Gradients for NBSR with dispersion

In an alternative modeling approach, we propose to model the dispersion parameters in relation to the underlying abundance level $\pi_{i,j}$. The gradients flow back to beta from the loss function (log posterior) to the dispersion parameters $\phi$ and mean parameters $\mu$ through $\pi$. The gradient for $\log p(\beta)$ is unchanged so we show here the gradient for the log likelihood. Note that under this model, we have dispersion for each sample-feature, $\phi_{ij}$ and hence, Negative Binomial distribution is parameterized with $r_{ij} = 1/\phi_{ij}$:

$$
\begin{aligned}
\log p(y_{i,j}|\beta) =& r_{i,j} \log p_{i,j} + y_{ij} \log(1 - p_{ij}) + && (26)\\
& \log \Gamma(y_{ij} + r_{ij}) - \log \Gamma(r_{ij}) - \log \Gamma(y_{ij}), && (27)
\end{aligned}
$$

The gradient is given by:

$$
\begin{aligned}
\nabla_{k,d} \log p(y_{i,j}|\beta) =& r_{i,j} \nabla_{k,d} \log p_{i,j} + y_{ij} \nabla_{k,d} \log(1 - p_{ij}) + && (28)\\
& \nabla_{k,d} r_{i,j} (\psi(y_{ij} + r_{ij}) - \psi(r_{ij}) + \log p_{i,j}), && (29)
\end{aligned}
$$

where $\psi$ denotes the Digamma function. First, we derive $\nabla_{k,d} r_{ij}$.

$$\nabla_{k,d} r_{ij} = -\frac{\nabla_{k,d} \phi_{ij}}{\phi_{ij}^2}, \tag{30}$$

where $\phi_{ij} = \exp(b_0 + b_1 \log \pi_{ij} + w' z_i)$.

$$
\begin{aligned}
\nabla_{k,d} \phi_{ij} &= \phi_{ij} \left( b_1 \frac{\nabla_{k,d} \pi_{ij}}{\pi_{ij}} \right) && (31)\\
&= \phi_{ij} b_1 x_{i,d} (1[j = k] - \pi_{ik}). && (32)
\end{aligned}
$$

Plug-in above, we get,

$$\nabla_{k,d} r_{ij} = -\frac{b_1 x_{i,d}(1[j = k] - \pi_{ik})}{\phi_{ij}}. \tag{33}$$

Therefore, Eq 29 is given by,

$$\frac{-b_1 x_{i,d}(1[j = k] - \pi_{ik})}{\phi_{ij}} (\psi(y_{ij} + r_{ij}) - \psi(r_{ij}) + \log p_{i,j}). \tag{34}$$

To derive Eq 28, we first express Eq 26 (note: $1 - p_{ij} = \phi_{ij} \mu_{ij}^2 / \sigma_{ij}^2$):

$$r_{i,j} \log p_{i,j} + y_{ij} \log(1 - p_{ij}) = r_{i,j}(\log \mu_{ij} - \log \sigma_{ij}^2) + y_{ij}(\log \phi_{ij} + \log \mu_{ij}^2 - \log \sigma_{ij}^2). \tag{35}$$

Now the gradient of $\sigma^2_{ij}$,

$$\begin{aligned}
\nabla_{k,d}\sigma^2_{i,j} &= \nabla_{k,d}\mu_{i,j} + \phi_{ij}\nabla_{k,d}\mu^2_{i,j} + \mu^2_{i,j}\nabla_{k,d}\phi_{ij} && (36)\\
&= x_{i,d}(1[j=k]-\pi_{i,k})(\mu_{i,j} + 2\phi_{ij}\mu^2_{i,j} + b_1\phi_{ij}\mu^2_{ij}) && (37)\\
&= x_{i,d}(1[j=k]-\pi_{i,k})(\sigma^2_{i,j} + \phi_{ij}\mu^2_{i,j}(1+b_1)) && (38)\\
&= x_{i,d}(1[j=k]-\pi_{i,k})(\sigma^2_{i,j} + (\sigma^2_{ij}-\mu_{i,j})(1+b_1)). && (39)
\end{aligned}$$

Hence, the gradient of $\log\sigma^2_{ij}$ is given by,

$$\nabla_{k,d}\log\sigma^2_{i,j} = x_{i,d}(1[j=k]-\pi_{i,k})(1+(1-p_{ij})(1+b_1)). \tag{40}$$

The gradient of Eq 28 is given by,

$$r_{ij}x_{id}(1[j=k]-\pi_{ik})(1-p_{ij})(1+b_1) + y_{ij}x_{id}(1[j=k]-\pi_{ik})(p_{ij}(1+b_1)). \tag{41}$$

$$\begin{aligned}
r_{ij}\nabla_{k,d}\log p_{i,j} &= r_{ij}(\nabla_{k,d}\log\mu_{ij} - \nabla_{k,d}\log\sigma^2_{ij}) && (42)\\
&= r_{ij}\left(\frac{\nabla_{k,d}\mu_{ij}}{\mu_{ij}} - \frac{\nabla_{k,d}\sigma^2_{ij}}{\sigma^2_{ij}}\right) && (43)\\
&= r_{ij}\left(\frac{x_{id}\mu_{ij}(1[j=k]-\pi_{ik})}{\mu_{ij}} - \frac{\nabla_{k,d}\sigma^2_{ij}}{\sigma^2_{ij}}\right) && (44)\\
&= r_{ij}\left(x_{id}(1[j=k]-\pi_{ik}) - \frac{x_{id}(1[j=k]-\pi_{ik})(\sigma^2_{ij}+\phi_{ij}\mu^2_{ij}(1+b_1))}{\sigma^2_{ij}}\right) && (45)\\
&= r_{ij}\left(x_{id}(1[j=k]-\pi_{ik}) - x_{id}(1[j=k]-\pi_{ik})(1+(1-p_{ij})(1+b_1))\right) && (46)\\
&= r_{ij}x_{id}(1[j=k]-\pi_{ik})(1-(1+(1-p_{ij})(1+b_1))) && (47)\\
&= -r_{ij}x_{id}(1[j=k]-\pi_{ik})(1-p_{ij})(1+b_1) && (48)
\end{aligned}$$

$$\begin{aligned}
y_{ij}\nabla_{k,d}\log(1-p_{i,j}) &= y_{ij}(\nabla_{k,d}\log\phi_{ij} + \nabla_{k,d}\log\mu^2_{ij} - \nabla_{k,d}\log\sigma^2_{ij}) && (49)\\
&= y_{ij}\left(\frac{\nabla_{k,d}\phi_{ij}}{\phi_{ij}} + \frac{\nabla_{k,d}\mu_{ij}}{\mu^2_{ij}} - \frac{\nabla_{k,d}\sigma^2_{ij}}{\sigma^2_{ij}}\right) && (50)\\
&= y_{ij}\left(\frac{\phi_{ij}b_1x_{id}(1[j=k]-\pi_{ik})}{\phi_{ij}} + \frac{2x_{id}\mu^2_{ij}(1[j=k]-\pi_{ik})}{\mu^2_{ij}} - \right. && (51)\\
&\qquad\left.\frac{x_{id}(1[j=k]-\pi_{ik})(\sigma^2_{ij}+\phi_{ij}\mu^2_{ij}(1+b_1))}{\sigma^2_{ij}}\right) && (52)\\
&= y_{ij}\left(b_1x_{id}(1[j=k]-\pi_{ik}) + 2x_{id}(1[j=k]-\pi_{ik}) - \right. && (53)\\
&\qquad\left. x_{id}(1[j=k]-\pi_{ik})(1+(1-p_{ij})(1+b_1))\right) && (54)\\
&= y_{ij}x_{id}(1[j=k]-\pi_{ik})(b_1+2-(1+(1-p_{ij})(1+b_1))) && (55)\\
&= y_{ij}x_{id}(1[j=k]-\pi_{ik})(1+b_1)p_{ij}. && (56)
\end{aligned}$$

### 4 Inference over log relative risk

Let $w \in \Omega$ be a categorical variable indicating experimental condition. We are interested in the relative change in the feature $j$ across two or more experimental conditions. Let us denote by $\mathbf{x}_i = (\mathbf{w}_i, \mathbf{z}_i)$ where $\mathbf{w}_i$ is a one-hot encoding denoting the experimental condition for sample $i$. We further decompose the model parameters by $\beta_j = (\theta_j, \gamma_j)$ where $\theta_j$ denotes the parameters corresponding to $\mathbf{w}_i$ and $\gamma_j$ for $\mathbf{z}_i$. Consider a multivariate function $g : \boldsymbol{\beta} \to \mathbb{R}^J$ with $g_j(\boldsymbol{\beta}) = \log RR_{j|\mathbf{z}}(\boldsymbol{\beta})$, where

$$\log RR_{j|\mathbf{z}} = \log \frac{\pi_{j|w_1,\mathbf{z}}}{\pi_{j|w_0,\mathbf{z}}}, \tag{57}$$

with $w_0, w_1 \in \Omega$ denoting the two experimental conditions. We can derive standard errors for $g_j$ by relying on the delta method and asymptotic normality of the MAP estimate $\boldsymbol{\beta}$, see for example, (Agresti, 2012, Chapter 3):

$$\nabla g(\boldsymbol{\beta})^T \Sigma \nabla g(\boldsymbol{\beta}).$$

Similarly to above, we provide the derivative of $g_j$ w.r.t. $\beta_{k,d}$.

$$\log \frac{\pi_{j|w_1,\mathbf{z}}}{\pi_{j|w_0,\mathbf{z}}} = \log \pi_{j|w_1,\mathbf{z}} - \log \pi_{j|w_0,\mathbf{z}}. \tag{58}$$

Using the gradient for $\pi_j$ derived above:

$$\log \frac{\pi_{j|w_1,\mathbf{z}}}{\pi_{j|w_0,\mathbf{z}}} = z_{w_1,d}(1[j=k] - \pi_{j|w_1,\mathbf{z}}) - z_{w_0,d}(1[j=k] - \pi_{j|w_0,\mathbf{z}}). \tag{59}$$

17 # 5 Dirichlet parameter estimation

We use mean-precision representation of Dirichlet and the estimation procedure outlined in (Minka, 2000). The log probability density for dataset $D = (\boldsymbol{p}_n)$ is given by,

$$\log p(D|\boldsymbol{\alpha}) = N \log \Gamma(s) - N \sum_k \log \Gamma(s m_k) + N \sum_k (s m_k - 1) \log \bar{p}_k, \tag{60}$$

18 where $\log \bar{p}_k = \frac{1}{N} \sum_n \log p_{n,j}$ denotes the sufficient statistics.

The fixed-point iteration for $s$ given by,

$$\frac{1}{s_{new}} = \frac{1}{s} + \frac{1}{s} \left( \frac{d^2}{ds^2} \log p(D|s) \right)^{-1} \left( \frac{d}{ds} \log p(D|s) \right). \tag{61}$$

The fixed-point iteration for $\alpha_k$ is given by,

$$\Psi(\alpha_{k,new}) = \log \bar{p}_k - \sum_j m_j (\log \bar{p}_j - \Psi(s m_j)), \tag{62}$$

followed by setting

$$m_{k,new} = \frac{\alpha_{k,new}}{\sum_j \alpha_{j,new}}. \tag{63}$$

The fixed-point iteration for $\alpha_k$ requires inverting $\Psi$, which can be posed as finding the root of:

$$F(x) = \Psi(x) - \log \bar{p}_k + \sum_j m_j (\log \bar{p}_j - \Psi(s m_j)) = 0. \tag{64}$$

This can be accomplished using Newton's method:

$$x_{new} = x_{old} - \frac{F(x_{old})}{F'(x_{old})} = x_{old} - \frac{F(x_{old})}{\Psi'(x_{old})}. \tag{65}$$

We initialize $\mathbf{m}_0 = \frac{1}{N} \sum_n \boldsymbol{p}_n$ and use Stirling's approximation for $s$:

$$s_0 = \frac{(K-1)/2}{-\sum_k m_k \log \bar{p}_k / m_k}. \tag{66}$$

Table 1: The number of microRNA samples for each cell type used in the miRNA versus mRNA comparison analysis.

| Cell type | N |
|---|---|
| Fibroblast | 45 |
| Smooth muscle | 43 |
| Endothelial | 43 |
| Monocyte | 51 |
| T lymphocyte CD8 | 48 |
| T lymphocyte CD4 | 48 |
| B lymphocyte CD19 | 42 |
| Natural Killer Cells CD56 | 44 |

Table 2: The number of messenger RNA samples for each cell type used in the miRNA versus mRNA comparison analysis.

| Cell type | N |
|---|---|
| Cardiac Muscle Cell | 33 |
| Endothelial Cell | 81 |
| Lymphocytes | 685 |
| Fibroblast | 13 |
| Macrophage | 318 |
| Fat Cell | 74 |
| Smooth Muscle Cell | 15 |

Table 3: Sample counts used for estimating Dirichlet-Multinomial parameters for simulated data analysis.

| Cell type | N |
|---|---|
| B lymphocyte CD19 | 39 |
| Monocyte | 43 |
| Natural killer cells CD56 | 41 |
| Neutrophil | 20 |
| Red blood cell | 25 |
| T lymphocyte CD4 | 42 |
| T lymphocyte CD8 | 41 |

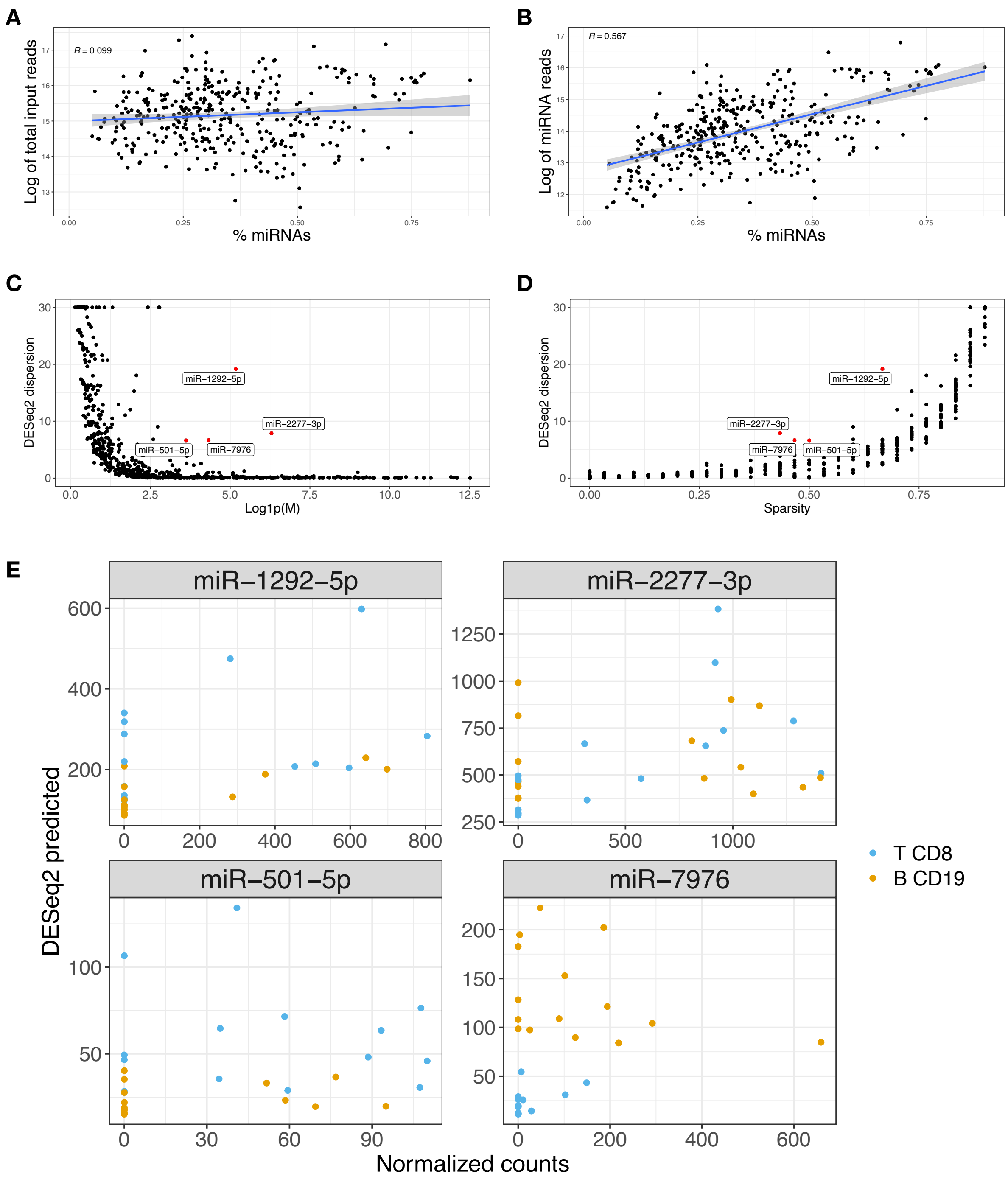
A
R = 0.099
Log of total input reads
% miRNAs
B
R = 0.567
Log of miRNA reads
% miRNAs
C
DESeq2 dispersion
miR−1292−5p
miR−2277−3p
miR−501−5p
miR−7976
Log1p(M)
D
DESeq2 dispersion
miR−1292−5p
miR−2277−3p
miR−7976
miR−501−5p
Sparsity
E
miR−1292−5p
miR−2277−3p
miR−501−5p
miR−7976
DESeq2 predicted
Normalized counts
T CD8
B CD19

Figure S1: **A**. Scatter plot of the total number of input reads versus percentage of miRNAs for 364 cell type samples from microRNAome dataset. Pearson correlation of 0.099 indicates that there is not a strong relationship. **B**. Linear correlation is observed between the number of miRNA reads versus percentage of miRNAs; Pearson correlation of 0.567 indicates that there is substantial variability in the miRNA capture rate. **C**. Scatter plot of $M$, the mean of normalized counts for miRNA $j$, in log scale versus dispersion estimation for $j$ from DESeq2 for T (CD8) vs B (CD19) dataset. The lowly expressed features tend to have high dispersion estimates. The miRNAs with relatively high expression and dispersion ($\log(M+1) > 3$ and DESeq2 dispersion $> 5$) are highlighted in red. **D**. Scatter plot of sparsity for feature $j$ versus DESeq2 dispersion estimation for $j$. The sparsity is measured by the number of samples with zero expression for $j$ divided by the total number of samples. All four of the miRNAs highlighted exhibit substantial sparsity. **E** Scatter plots of DESeq2 predicted expression level versus normalized counts using DESeq2's size factor estimation for miRNAs highlighted in red in **C**. The normalized counts are seen to be overdispersed leading to DESeq2 to produce large dispersion estimates to account for zero expression.

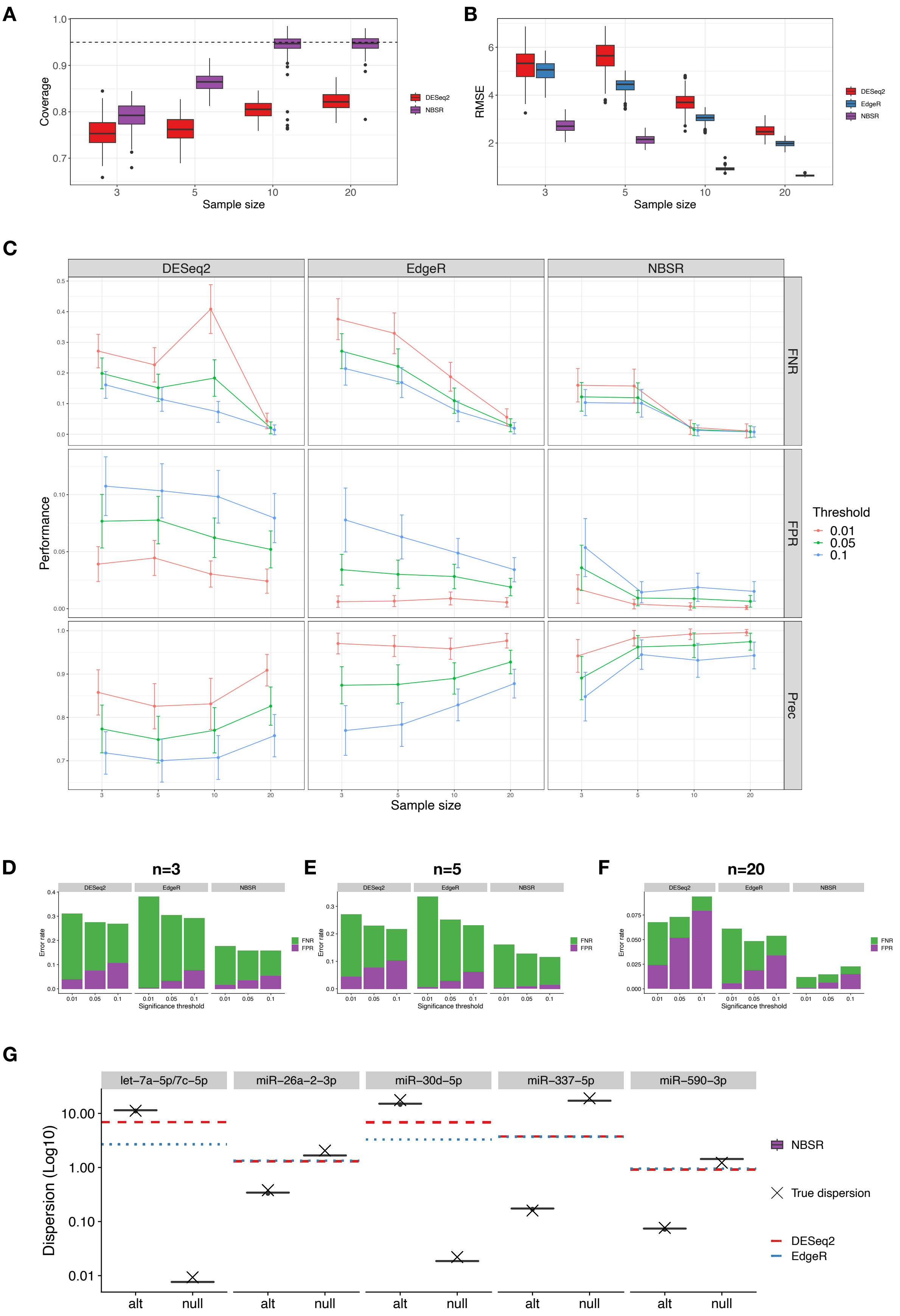

A
Coverage
Sample size
DESeq2
NBSR
B
RMSE
Sample size
DESeq2
EdgeR
NBSR
C
DESeq2
EdgeR
NBSR
FNR
FPR
Prec
Performance
Sample size
Threshold
0.01
0.05
0.1
D
n=3
E
n=5
F
n=20
Error rate
Significance threshold
FNR
FPR
G
let−7a−5p/7c−5p
miR−26a−2−3p
miR−30d−5p
miR−337−5p
miR−590−3p
Dispersion (Log10)
alt
null
NBSR
True dispersion
DESeq2
EdgeR

Figure S2: **A-B**. Confidence interval coverage of the true fold change and root mean squared error of the predicted fold change to the true fold change as the sample size (x-axis) is increased. EdgeR does not output confidence interval and hence, omitted from the figure. NBSR for samples $n \in \{3, 5\}$ uses Empirical Bayes estimation while for $n \in \{10, 20\}$ uses linear dispersion model. Boxplots depict IQR1-3. **C**. Precision, false positive rates, and false negative rates for three methods as the sample sizes and significance thresholds are varied. The confidence band indicates $\pm\sigma$ obtained over 20 replicates for each of the 7 parameter settings (total of 140). NBSR attains lowest error rate and highest precision at conservative threshold of 0.01. **D-F** Detailed breakdown of false positive and negative error rates for three methods at different threshold values. **G**. Dispersion estimation for a subset of the perturbed miRNAs for simulated data analysis with $n = 20$.

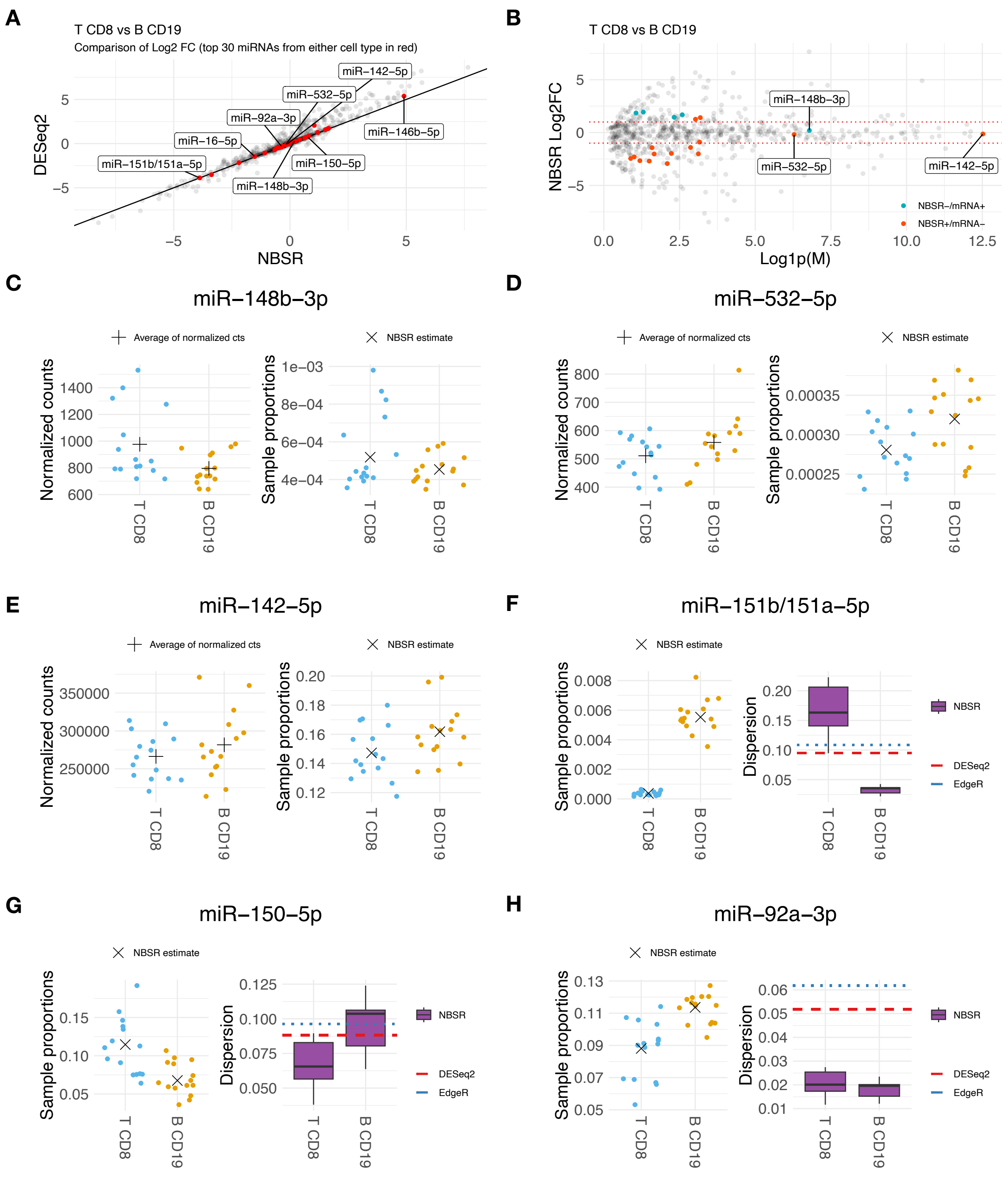
A
T CD8 vs B CD19
Comparison of Log2 FC (top 30 miRNAs from either cell type in red)
DESeq2
NBSR
miR−142−5p
miR−532−5p
miR−92a−3p
miR−16−5p
miR−146b−5p
miR−151b/151a−5p
miR−150−5p
miR−148b−3p
B
T CD8 vs B CD19
NBSR Log2FC
Log1p(M)
miR−148b−3p
miR−532−5p
miR−142−5p
NBSR−/mRNA+
NBSR+/mRNA−
C
miR−148b−3p
Average of normalized cts
NBSR estimate
Normalized counts
Sample proportions
T CD8
B CD19
D
miR−532−5p
E
miR−142−5p
F
miR−151b/151a−5p
Dispersion
NBSR
DESeq2
EdgeR
G
miR−150−5p
H
miR−92a−3p

Figure S3: **A**. Scatter plot of log2 fold change estimates between DESeq2 and NBSR for miRNAs used in the analysis of T CD8 versus B CD19 cells. Majority of the fold change estimates concur between the two methods. The highlighted in red are examined in more detail. **B**. MA-plot to highlight the differences between DESeq2 and NBSR. The NBSR+/mRNA- indicate miRNAs at FDR $< 0.05$ for NBSR but FDR $> 0.1$ for both DESeq2 and EdgeR while NBSR-/mRNA+ indicate miRNS at FDR $< 0.05$ for both DESeq2 and EdgeR while FDR $> 0.1$ for NBSR. Most of the disagreements are lowly expressed with the exception of miR-148b-3p (NBSR-/mRNA+) and miR-532-5p/miR-142-5p (NBSR+/mRNA-). **C-E**. Sample proportions, NBSR estimated proportion (marked by cross), and NBSR dispersion estimates for three miRNAs where mRNA methods and NBSR differed. **F**. In-depth look into a miRNA with high fold change due to low baseline. **G-H**. In-depth look into miRNAs with low fold change but large absolute difference in NBSR estimated proportion.

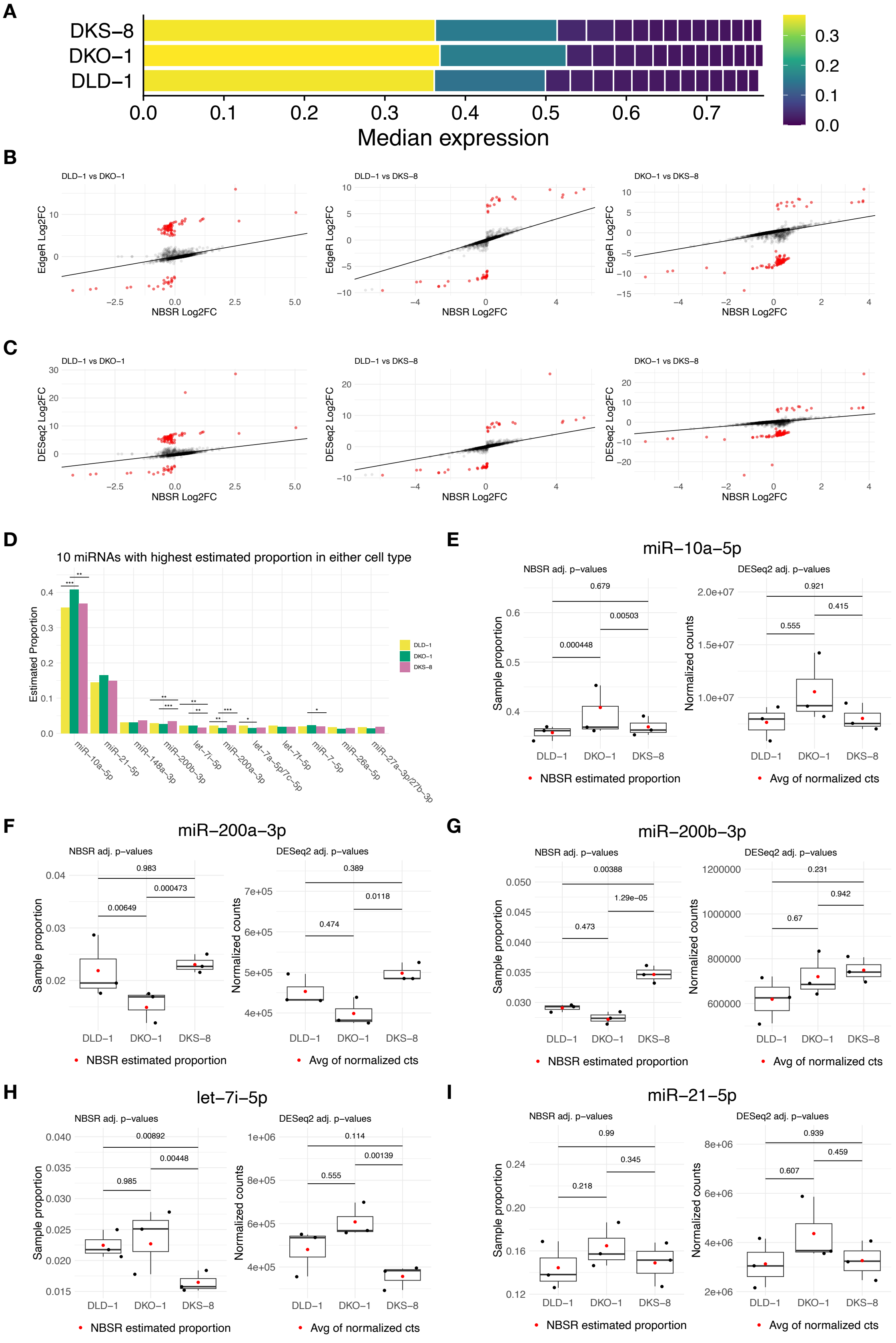

A
DKS−8
DKO−1
DLD−1
Median expression
B
DLD−1 vs DKO−1
DLD−1 vs DKS−8
DKO−1 vs DKS−8
EdgeR Log2FC
NBSR Log2FC
C
DLD−1 vs DKO−1
DLD−1 vs DKS−8
DKO−1 vs DKS−8
DESeq2 Log2FC
NBSR Log2FC
D
10 miRNAs with highest estimated proportion in either cell type
Estimated Proportion
DLD−1
DKO−1
DKS−8
miR−10a−5p
miR−21−5p
miR−148a−3p
miR−200b−3p
let−7i−5p
miR−200a−3p
let−7a−5p/7c−5p
let−7f−5p
miR−7−5p
miR−26a−5p
miR−27a−3p/27b−3p
E
miR−10a−5p
NBSR adj. p−values
0.679
0.00503
0.000448
Sample proportion
DESeq2 adj. p−values
0.921
0.415
0.555
Normalized counts
NBSR estimated proportion
Avg of normalized cts
F
miR−200a−3p
NBSR adj. p−values
0.983
0.000473
0.00649
Sample proportion
DESeq2 adj. p−values
0.389
0.0118
0.474
Normalized counts
NBSR estimated proportion
Avg of normalized cts
G
miR−200b−3p
NBSR adj. p−values
0.00388
1.29e−05
0.473
Sample proportion
DESeq2 adj. p−values
0.231
0.942
0.67
Normalized counts
NBSR estimated proportion
Avg of normalized cts
H
let−7i−5p
NBSR adj. p−values
0.00892
0.00448
0.985
Sample proportion
DESeq2 adj. p−values
0.114
0.00139
0.555
Normalized counts
NBSR estimated proportion
Avg of normalized cts
I
miR−21−5p
NBSR adj. p−values
0.99
0.345
0.218
Sample proportion
DESeq2 adj. p−values
0.939
0.459
0.607
Normalized counts
NBSR estimated proportion
Avg of normalized cts

Figure S4: **A**. Plot of the top 15 most highly expressed miRNAs in colon adenocarcinoma cell line data. The first two miRNAs account for close to 60% of the read counts. **B**. Comparison of log2 fold change estimates between EdgeR and NBSR. **C** Comparison of log2 fold change estimates between DESeq2 and NBSR. Both the DESeq2 and EdgeR produce large fold change estimates as highlighted in red. **D**. Top 10 miRNAs from either of the three cell lines along with the significance levels indicated by $***$ for $< 0.001$, $**$ for $< 0.01$, and $*$ for $< 0.05$. **E-I**. Side-by-side plots of sample proportion and NBSR estimated proportion (marked in red) and DESeq2 normalized counts and average of normalized counts (marked in red), along with BH adjusted p-values from respective methods. NBSR identifies which of the samples are significantly differential in the expression of miR-10a-5p, miR-200a-3p, miR-200b-3p, and let-7i-5p. NBSR and DESeq2 agrees on miR-21-5p. Boxplots depict IQR 1-3.
